# Fusion of a functional glutaredoxin to the radical-generating subunit of ribonucleotide reductase

**DOI:** 10.1101/373563

**Authors:** Inna Rozman Grinberg, Daniel Lundin, Margareta Sahlin, Mikael Crona, Gustav Berggren, Anders Hofer, Britt-Marie Sjöberg

**Affiliations:** From the Department of Biochemistry and Biophysics, Stockholm University Sweden; From the Swedish Orphan Biovitrum AB, Stockholm, Sweden; From the Department of Chemistry, Uppsala University Uppsala, Sweden; From the Department of Medical Biochemistry and Biophysics, Umeå University Umeå, Sweden

**Author notes:** To whom correspondence should be addressed: Britt-Marie Sjöberg, Department of Biochemistry and Biophysics, Stockholm University, SE-10691 Stockholm, Sweden.

**Keywords:** allosteric regulation, ATP-cone, dATP inhibition, dimanganese, dithiol, monothiol mechanism, tetramers, radical mechanism, redox activity, ribonucleotide reductase

## Abstract

Class I ribonucleotide reductase (RNR) consists of a catalytic subunit (NrdA) and a radical-generating subunit (NrdB) that together catalyse reduction of the four ribonucleotides to their corresponding deoxyribonucleotides. *Facklamia ignava* NrdB is an unprecedented fusion protein with N-terminal add-ons of a glutaredoxin (Grx) domain followed by an ATP-cone. Grx, which in general is encoded elsewhere in the genome than is the RNR operon, is a known physiological reductant of RNRs. Here we show that the fused Grx domain functions as an efficient reductant of the *F. ignava* class I RNR via the common dithiol mechanism and interestingly also via a monothiol mechanism, although less efficiently. A Grx that utilizes either or of these two reaction mechanisms has to our knowledge not been observed with a native substrate before. The ATP-cone, which is commonly found as an N-terminal domain of the catalytic subunit of RNRs, is an allosteric on/off switch that promotes dNDP reduction in presence of ATP and inhibits the enzyme activity in presence of dATP. Here we show that dATP bound to the ATP-cone of *F. ignava* NrdB promotes formation of tetramers that are unable to form enzymatically competent complexes with *F. ignava* NrdA. The ATP-cone binds two molecules of dATP, but only one molecule of the activating nucleotide ATP. *F. ignava* NrdB contains the recently identified radical factor Mn_2_^III/IV^. We show that NrdA from the firmicute *F. ignava* can form a catalytically competent RNR with the Mn_2_^III/IV^-containing NrdB from the flavobacterium *Leeuwenhoekiella blandensis*.

## Introduction

Ribonucleotide reductase (RNR) is an essential enzyme that catalyze the synthesis of the DNA building blocks (dNTPs) by reduction of the four ribonucleotides. RNR plays a key role in DNA synthesis and DNA repair, and consequently attracts biomedical interest as a potential target for antibacterial substances and for anticancer therapies. Currently, the RNR enzyme family comprises three different RNR classes and several subclasses. The three classes have a common reaction mechanism that builds on radical chemistry, but differ in the way they initiate the radical mechanism (1-5). The class I RNRs consist of a larger catalytic component (NrdA) and a smaller radical-generating metal-containing component (NrdB) where the dinuclear metal site differs between subclasses. Currently, class I RNRs have been subclassified based on radical cofactor type (subclasses Ia, Ib, Ic and Id) or evolutionary history (subclasses NrdA/B followed by a small letter plus subclass NrdE/F) (1,6). Metal content does not always follow phylogeny as two unrelated Mn_2_ subclasses exist, where one subclass contains a tyrosyl radical in the vicinity of a Mn^III^/Mn^III^ center (Ib, NrdE/F), and another recently identified subclass (Id, NrdAi/Bi) contains a mixed valent Mn^IV^/Mn^III^ metal center that harbors the unpaired electron (7-9). Moreover, in eukaryotic RNRs and several evolutionarily unrelated bacterial class I subclasses, the NrdB component contains a stable tyrosyl radical in the vicinity of a diferric metal center (Ia). In another bacterial subclass (Ic), a mutational change in the radical-carrying tyrosine to phenylalanine is accompanied by a mixed valent Mn^IV^/Fe^III^ metal center (10,11). Recently, a metal independent subclass (Ie) with an intrinsically modified dopa radical cofactor was discovered (12).

All class I RNRs contain a C-terminal redox-active cysteine pair in NrdA that functions as a reductant of a cysteine pair in the active site that is oxidized during catalysis. Physiological regeneration of active NrdA is performed by members of the redoxin family, with NADPH as ultimate electron source (13-15). Three types of redoxin have been found to reduce the C-terminal cysteines in class I RNRs: *i)* thioredoxin that receives the electrons from NADPH via thioredoxin reductase, *ii)* glutaredoxin (Grx) that receives the electrons from NADPH via glutathione reductase and glutathione (GSH), and *iii)* NrdH-redoxin that also receives the electrons from NADPH via thioredoxin reductase even though NrdH is more similar to Grx than to thioredoxin. Whereas the *nrdA* and *nrdB* genes are mostly encoded by an operon in bacteria, the *trx* and *grx* genes are usually found elsewhere in the genome. Only the *nrdH* gene is predominantly encoded by the same operon as the corresponding RNR genes, which for historical reasons in this particular subclass are called *nrdE* (encoding the catalytic subunit) and *nrdF* (encoding the radical-generating subunit).

We discovered an intriguing fusion of a *grx* gene to the the *nrdB* gene in the bacterium *Facklamia ignava*, resulting in an open reading frame encoding a fusion protein. The *F. ignava* NrdB fusion protein consists of an N-terminal Grx domain followed by an ATP-cone domain and then the radical-generating subunit. An N-terminal *grx* fusion to the *nrdB* gene in *Francisella tularensis* was noticed by us previously (16). The redoxin domain in both these fusion are most closely related to the *grxC* domain family (COG0695). Whereas the γ- proteobacterium *Francisella tularensis* is a well studied human pathogen causing tularemia (17), the Firmicutes genus *Facklamia* was first described in 1997 and has since been identified in samples from a wide range of animals and as a human pathogen (18-20).

RNR has been described as a textbook example of allosteric regulation in enzymes, and employs two different allosteric mechanisms to regulate the synthesis of dNTPs (21). One common mechanism regulates the balance between the four dNTPs in a sophisticated feedback control at the specificity site (s-site). An additional allosteric regulation occurs in a separate domain called the ATP-cone and works as a general on/off switch, the overall activity site (a-site). In simple terms, the enzyme is active when ATP is bound and when dATP is bound the enzyme is turned off. We have recently shown that the ATP-cone can be horizontally transferred between different RNRs and even to different subunits of the holoenzyme (8,22). In an overwhelming number of cases the ATP-cone is an N-terminal domain of the catalytic subunit of RNR (22). *F. ignava* RNR instead carries an ATP-cone in its NrdB protein, between the N-terminal Grx domain and the radical-generating domain. We have recently reported a similar N-terminal ATP-cone fusion to NrdB in *Leewenhoekiella blandensis* (8). Both these fusion proteins belong to the NrdBi subclass, which harbors a few additional ATP-cone::NrdB fusions.

In this study we have used the *F. ignava* RNR to study two major questions: does the fused Grx domain function as a reductant for the holoenzyme, and does the fused ATP-cone function as a general on/off switch? To investigate this, we used a series of biochemical assays to show that the Grx domain is indeed an efficient reductant of *F. ignava* RNR, and that the fused ATP-cone domain is a functional allosteric domain.

## Results

### Glutaredoxin-fusions to RNR components

The 496-residue *F. ignava* (Firmicutes) NrdB fusion protein consists of an N-terminal Grx domain (residues 4-61, with the characteristic cysteine pair at residues 12 and 15) followed by an ATP-cone domain (residues 84-169) and thereafter the NrdB proper (cf Fig. S5). The *nrdA* gene is located 46 nucleotides downstream of the *nrdB* gene, and the two genes conceivably form an operon (Fig. 1). The *F. ignava* NrdB is a member of the NrdBi phylogenetic subclass (http://rnrdb.pfitmap.org), like all other NrdBs in which we have detected N-terminal ATP-cones (8).

**Figure 1.**
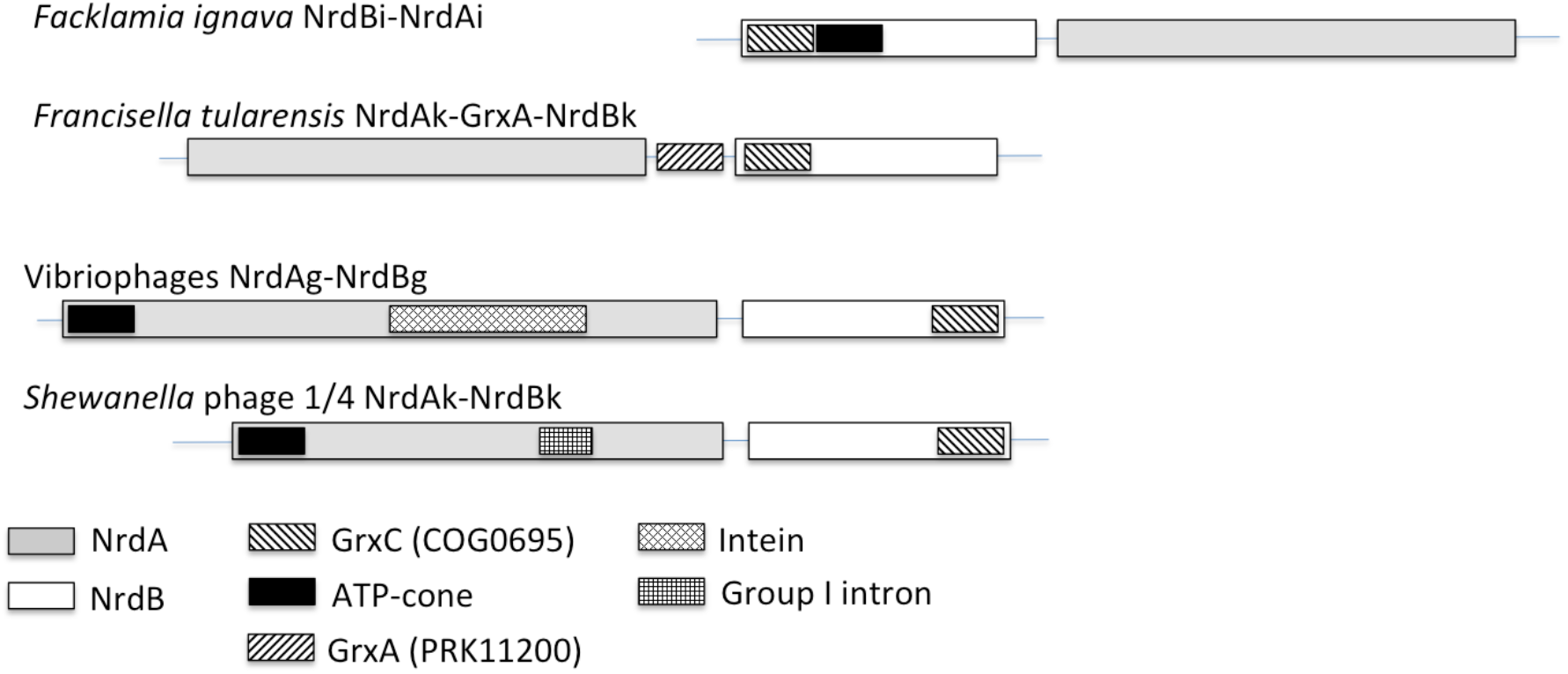
The *F. ignava nrdAB* operon and some other class I RNR operons with *grx* fusions. Transcriptional and translational directions are from left to right. The *Francisella tularensis* operon occur in all *Francisella* spp. and the displayed vibriophage operon is found in most vibriophages (Table S1).

Spurred by the discovery of the Grx fusion to *F. ignava* NrdBi, we performed a search of the RefSeq database for combinations of RNR proteins and Grx domains. Grx fusions were found in all RNR components (NrdA, NrdB, NrdD and NrdJ), in some cases together with an ATP-cone (Fig. 1, Table S1). Grx fused to NrdB were detected in all *Francisella* spp. and *Allofrancisella guangzhouensis* (both γ-proteobacteria; subclass NrdBk), and in 24 viruses (NrdBe, NrdBg and NrdBk) (Fig. 1, Table S1). In addition, a *grx::nrdE* fusion was found in *Streptococcus pneumoniae*, a *grx::nrdD* fusion in *Lachnospiraceae bacterium TWA4*, a *grx::nrdJ* fusion in *Labrenzia aggregata* and *grx::nrdA* fusions in 2 viruses (Table S1).

As many Firmicutes lack glutathione and instead produce another low molecular weight reductant called bacillithiol (23), we also searched the *F. ignava* genome for presence of GSH biosynthesis and bacillithiol biosynthesis genes (*gshA, gshB, gshF, bshA, bshB*1, *bshB*2, *bshC*). *F. ignava* and the other *Facklamia* spp. but one encode the bifunctional *gshF* gene that is primarily found in Firmicutes (24). The *Facklamia gshF* has extensive similarity primarily to the *gshA* gene (Table S2). The glutathione reductase gene *gor* was only found in *Facklamia sourekii*. The closest ortholog in *F. ignava* is a mercury(II) reductase and a dihydrolipoyl dehydrogenase, both with ca 50% similarity to *F. sourekii gor*. There were no genes corresponding to the bacillithiol biosynthesis genes in any *Facklamia* spp., apart from a glycosyl transferase gene with some similarity to *bshA*. Our results show that *F. ignava* and other *Facklamia* spp. have the capacity to synthesize GSH.

### Redox activity of the NrdB-fused glutaredoxin

Using a series of cysteine-to-serine mutant proteins we have delineated the reaction mechanism of the fused Grx domain. Grx proteins usually reduce RNRs via a dithiol mechanism, but e.g. a human Grx has been reported to work via a glutathionylation mechanism (25-27). To test the capacity of the Grx domain in *F. ignava* NrdB to perform a dithiol reduction, we constructed two mutant proteins with a serine instead of cysteine in one or the other of the two redox-active residues in the Grx domain (C12S and C15S) and the corresponding double mutant (C12S/C15S).

In a first set of experiments the mutants were compared to the wild type protein in a redox cycle with the artificial substrate 2-hydroxyethyl disulfide (HED). As evident from figure 2A, the wild type and C15S mutant proteins reduced the HED substrate, whereas the C12S mutant and the double mutant did not. The *K_m_* for HED was 0.6 ±0.09 mM for the wild type protein and 1.3 ±0.24 mM for the C15S protein and the *V_max_* was approximately 2-fold higher for the wild type compared to C15S at saturating HED (Fig. 2B). In a GSH titration experiment with constant HED, the K_m_ for GSH was 3 ±0.9 mM for the C15S mutant protein and the rate was 44 μmol/min (Fig. 2C) corresponding to a *k_cat_* of 1.5 s^−1^. Activity in the absence of one cysteine demonstrates that the Grx domain in *F. ignava* NrdB can work via a monothiol mechanism utilizing Cys12 as the redox-active cysteine in presence of HED. The behavior of the wild type protein in the GSH titration experiment (Fig. 2C) cannot be explained by a pure dithiol reaction mechanism. One possible explanation is that a monothiol mechanism may interfere at higher GSH concentrations.

**Figure 2.**
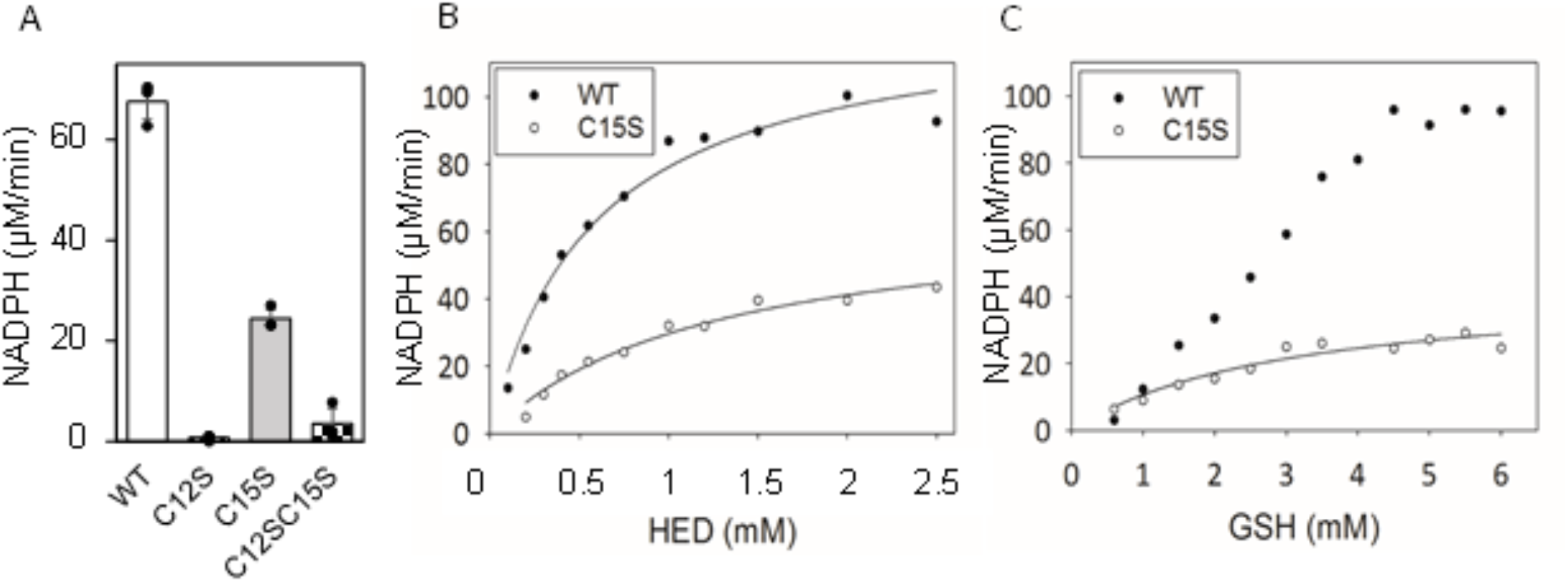
HED reduction capacity of *F. ignava* NrdB. A) 0.1 μM wild type and mutant proteins in the presence of 0.75 mM HED and 4 mM GSH. Assays were performed in triplicate with standard deviations shown. B) HED titration of wild type and C15S proteins in the presence of 4 mM GSH. C) GSH titration of wild type and C15S proteins in presence of 0.75 mM HED.

In a second set of experiments we compared the ability of the wild type and mutant Grx domains to function as reductants in RNR assays. High specific activity (*k_cat_* 1.4 ±0.06 s^−1^) with an apparent *K_m_* for GSH of 1.2 ±0.2 mM was only obtained with the wild type protein (Fig. 3). Of the mutant proteins, C12S and the C12S/C15S were deficient in ribonucleotide reduction both with 4 mM and 10 mM GSH, whereas their specific activity was on par with the wild type enzyme when the Grx domain was bypassed using DTT as reductant (Fig. 3C, inset). Interestingly, the C15S mutant promoted a low but significant GSH-dependent ribonucleotide reductase activity, as measured both as consumption of NADPH (Fig. 3A) and as formation of dCDP (Fig. 3C), but it was not possible to reach a *V_max_* for the RNR activity of the C15S protein even at 20 mM GSH (Fig. 3B and data not shown). The GSH concentration of *Facklamia* spp. is not known, but GSH concentrations in studied bacteria range between 0.1 and 10 mM, with Firmicutes generally on the high side (Masip, 2006 #224;Fahey, 2013 #183}. Conceivably, the Grx fused to *F. ignava* NrdB is most efficiently promoting turnover of the *F. ignava* RNR via a dithiol mechanism, and at 10 mM GSH concentration the C15S mutant protein can promote approximately 4-fold less efficient ribonucleotide reduction via a monothiol mechanism involving Cys12 (Fig. S1).

**Figure 3.**
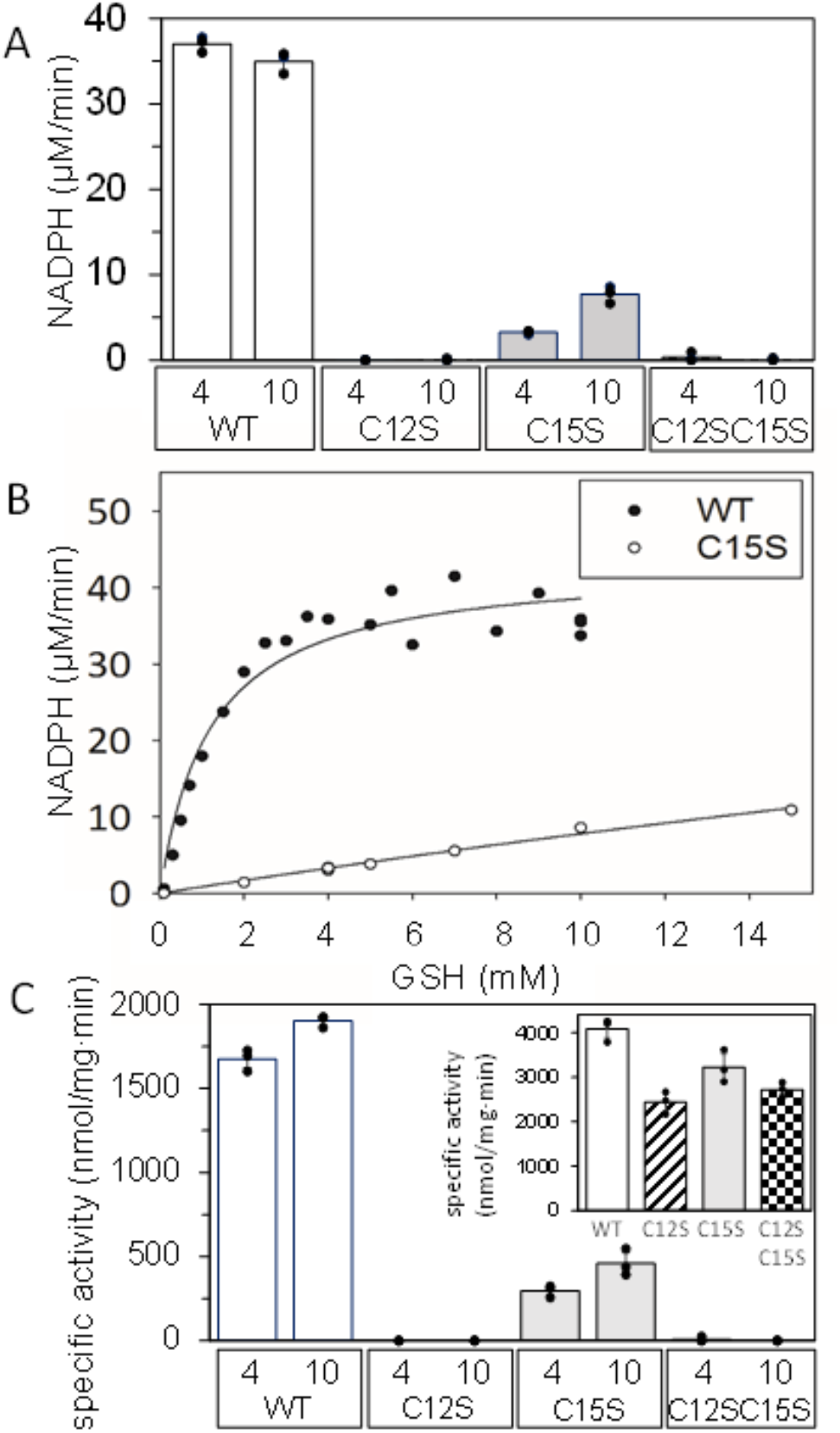
GSH-dependent RNR activity of *F. ignava* NrdB wildtype and mutant proteins. CDP was used as substrate and 3 mM ATP as effector. A) Enzyme activity measured as NADPH consumption in presence of 0.5 μM NrdB. B) GSH-dependent NADPH consumption of the wild type (•) and the C15S (○) NrdB. C) GSH-dependent specific activity measured as dCDP formation. *Inset:* DTT (10 mM) dependent specific activity measured as dCDP formation. GSH concentrations (4 and 10 mM) are indicated in A and C. Assays in A and C were performed in triplicate with standard deviations shown.

### Substrate specificity regulation of *F. ignava* RNR via the s-site

Using a four-substrate activity assay in the presence of saturating concentrations of the substrate specificity site (s-site) effectors ATP, dTTP or dGTP, we found that *F. ignava* RNR has a similar specificity regulation pattern to most characterized RNRs (3). ATP stimulated the reduction of CDP and UDP, whereas dTTP stimulated the reduction of GDP, and dGTP stimulated the reduction of ADP and GDP (Fig. 4). There is also a low activity of predominantly CDP reduction in the absence of allosteric effectors. Using mixtures of allosteric effectors, we observed that dTTP-induced GDP reduction increased in the presence of ATP (data not shown), as is commonly seen in RNRs (3).

**Figure 4.**
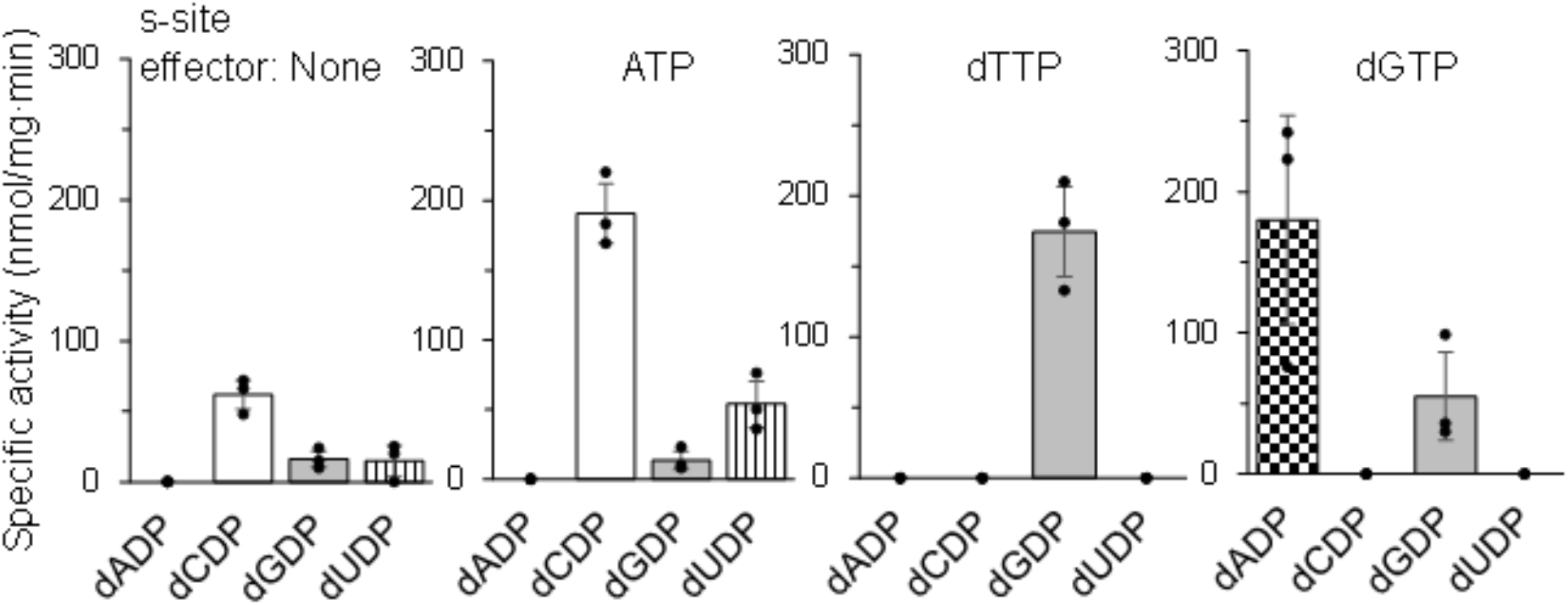
Substrate specificity of *F. ignava* class I RNR. Enzyme assays were performed in mixtures with 0.5 mM of each of the four substrates (ADP, CDP, GDP, UDP) and a saturating concentration of one effector nucleotide at a time. Assays were performed in triplicate with standard deviations shown.

### Overall activity of *F. ignava* RNR is regulated via the NrdB-linked ATP-cone

We performed a series of activity assays with CDP as substrate to elucidate the potential roles of ATP and dATP in activating and inhibiting the enzyme. The presence of ATP activated the enzyme, while dATP showed a dual effect and activated the enzyme when used at low concentrations and was inhibitory at 3 μM and higher (Fig. S2). The kinetics are complex, as ATP and dATP can bind both to the s-site in NrdA as well as the ATP-cone in NrdB. To analyze the effects of ATP or dATP at the ATP-cone of NrdB, the specificity site of NrdA was saturated with dTTP and GDP was used as substrate, giving a starting specific activity (normalized to 100% in Fig. 5) even in the absence of added ATP. In wild type NrdB *K_L_* for ATP-dependent activation was 47 ± 12 μM (Fig. 5A), and *K_i_* for dATP-dependent inhibition was 1.3 ± 0.23 μM (Fig. 5B). The activity of the deletion mutant that lacks both the Grx domain and the ATP-cone (NrdB∆169) was not affected by addition of either ATP or dATP (Fig. 5A,B). The initial *k_cat_* of the NrdB∆169 in the presence of dTTP-loaded NrdA was 2.2 s^−1^, i.e. almost 3 times higher than that of full-length NrdB (0.8 s^−1^), however ATP addition increased the activity of wild type NrdB to 1.8 s^−1^ (Fig. 5A); i.e. on par with the NrdB∆169 mutant and other RNR enzymes. Titration with dADP inhibited the wild type enzyme activity (Fig. S3), although less strongly than did dATP.

**Figure 5.**
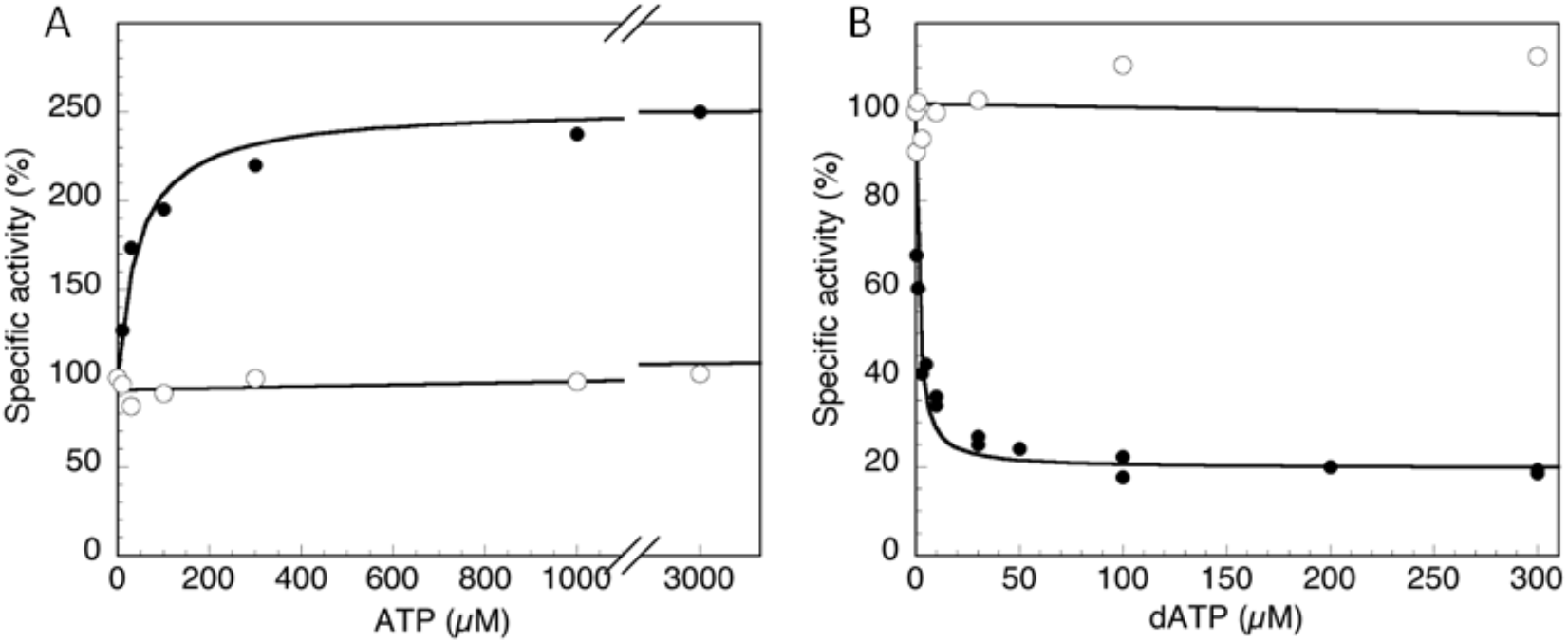
Inhibition and activation of wild type and mutant enzyme activity by dATP and ATP. A) ATP titration, and B) dATP titration of enzyme loaded with 2 mM dTTP and GDP as substrate. Specific activities of NrdB proteins were measured with a 10-fold excess of NrdA. Wild type NrdB (•) had a starting activity of 830 ±120 nmol/min•mg in the absence of added ATP and reached a *V_max_* of 1850 ±200 nmol/min•mg (*k_cat_* = 1.8 s^−1^) in presence of ATP, whereas NrdB∆169 (○) had a specific activity of 3400 ±500 nmol/min•mg (*k_cat_* = 2.2 s^−1^) in the absence of ATP that was not affected by addition of ATP or dATP.

### dATP binding to NrdB induces formation of higher oligomeric complexes

To elucidate the mechanism of allosteric overall activity regulation governed by the NrdB-linked ATP-cone, oligomer-distribution experiments were performed by gas-phase electrophoretic macromolecule analysis (GEMMA). GEMMA analysis showed that the NrdB subunit (β) is in a dimer-tetramer equilibrium and the tetramer formation is stimulated by dATP and suppressed by ATP (Fig. 6A). If the ATP-cone is removed as in NrdB∆169, the protein loses the ability to form tetramers, indicating that the process is dependent on the ATP-cone (Fig. 6B). In the Grx deletion mutant the ability to form tetramers is decreased but not lost completely (Fig. 6B). The NrdA subunit (α) is in a monomer-dimer equilibrium favoring dimers, especially in the presence of dATP where the monomers are below the detection limit (Fig. 6C). When both proteins were mixed together with dATP, an additional peak corresponding to an α_2_β_4_ complex appeared and to a minor extent also an α_4_β_4_ complex (Fig. 6D). In the absence of allosteric effectors or in the presence of ATP, α _2_β_2_ complexes were formed instead. The subunit compositions of the 234, 344 and 470 kDa peaks were determined by comparing the results with each subunit alone. NrdB tetramer formation is very inefficient in the absence of effectors or in the presence of ATP, indicating that the 234 peak only to a minor extent can be explained by NrdB tetramers and to most part contains α_2_β_2_ complexes, which is the result if the major two species NrdA and NrdB dimers interact with each other. In the presence of dATP, the two major species NrdA dimers and NrdB tetramers interact to form the α_2_β_4_ complex and to some extent also an α_4_β_4_ complex if an additional NrdA dimer binds.

**Figure 6.**
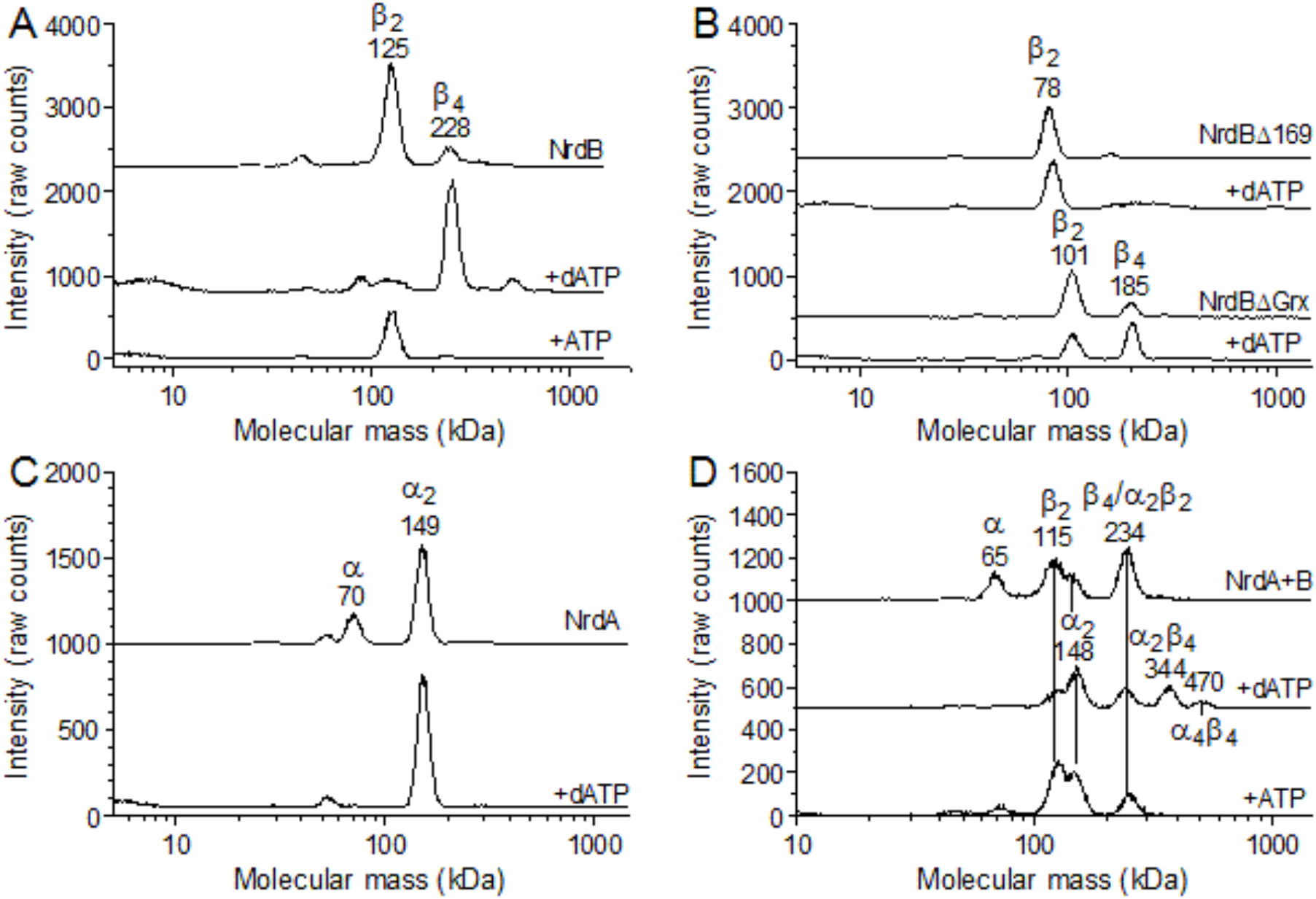
GEMMA analysis of the *F. ignava* ribonucleotide reductase. A) Analysis of 0.05 mg/ml NrdB (0.9 μM) in the absence or presence of 50 μM dATP or 100 μM ATP. B) Similar experiments as in A but with NrdB mutant proteins lacking the ATP cone (NrdB∆169) or the Grx domain (NrdB∆Grx) analyzed with and without 50 μM dATP. C) Analysis of 0.05 mg/ml NrdA protein in the absence or presence of 50 μM dATP. D) Experiments with NrdA-NrdB mixtures containing 0.025 mg/ml of each protein and no effector, 50 μM dATP or 100μM ATP.

To complement the GEMMA analyses of oligomer formation, we performed analytical size exclusion chromatography (Fig. 7) using higher protein concentrations and physiologically reasonable concentrations of effectors (3 mM ATP and 0.1 mM dATP) (28,29). The SEC experiments confirmed the GEMMA results. The NrdA protein was a dimer and this ability was enhanced by binding of dATP as well as ATP to its s-site (Fig. 7A). The NrdB subunit doubled in mass in the presence of dATP as compared to ATP, supporting the GEMMA result that it is a dimer with ATP and tetramer with dATP (Fig. 7B). In SEC, both the dimer and tetramer had larger masses than expected, indicating that the shape of the protein is not perfectly globular. Without effector, the NrdB protein seems to be a dimer that gradually goes through a transition to a larger species at higher protein concentration (Fig. 7B). This is in agreement with the GEMMA results that there is a dimer-tetramer equilibrium with the majority of the protein being dimeric (Fig. 6A). When the NrdA and NrdB proteins were mixed, they formed an α_2_β_2_ complex with ATP and without effector and a larger species with dATP. There was a gradual movement to a larger species when the protein concentration is increased up to a mass indicating an α_4_β_4_ complex.

**Figure 7.**
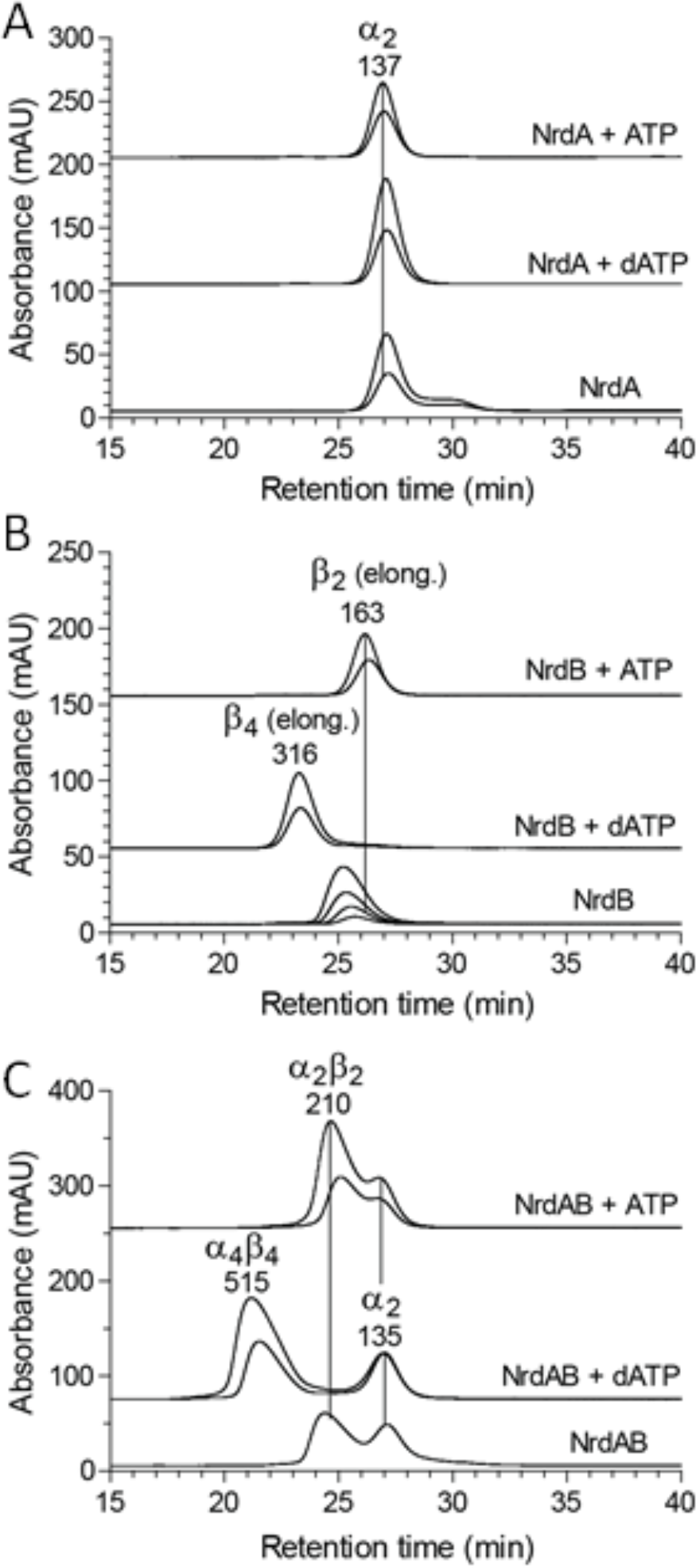
Size exclusion chromatography analysis of *F. ignava* RNR components in the presence of nucleotides. A) 5 and 10 μM of the NrdA subunit was analyzed in the presence of 3 mM ATP, 100 μM dATP or without effector. B) Corresponding analysis of the NrdB subunit. In this case the experiment without effector was performed at 1.25, 2.5, 5 and 10 μM protein. The position of the peaks indicate a larger size than expected, which is typical for elongated proteins, and the interpretation above the peaks is based on a comparison with the GEMMA results. C) Analysis of the combination of both subunits. Each subunit was used at 10 and 20 μM concentration except in the experiment without effector where only 10μM was used.

Binding of nucleotides to *F. ignava* NrdB was investigated using isothermal titration calorimetry (ITC). Binding curves for dATP and ATP to NrdB at 20°C were consistent with a single set of binding sites (Fig. 8). In dATP titrations the fitted apparent N value was significantly above one (N = 1.4 ±0.1), suggesting that the protein binds two dATP molecules per ATP-cone provided our preparation contains approximately 70% active protein. Fit of ATP titrations, performed with the same protein preparation and at the same day resulted in N = 0.55 ±0.02, suggesting binding of only one ATP per ATP-cone. *K_d_* for the different nucleotides (Fig. 8E) indicated a 20-fold lower affinity for ATP compared to dATP. Thermodynamic parameters (Fig. 8E) indicated that the interactions are predominantly enthalpy driven, with negative ΔH values of −80 and −60 kJ/mol for dATP and ATP, respectively. As earlier observed for *L. blandensis* NrdB (8) dADP also binds to the ATP-cone of *F. ignava* NrdB with a *K_d_* of 5.8 μM at 25°C, i.e. considerably weaker than the *K_d_* for dATP (Fig. S4).

**Figure 8.**
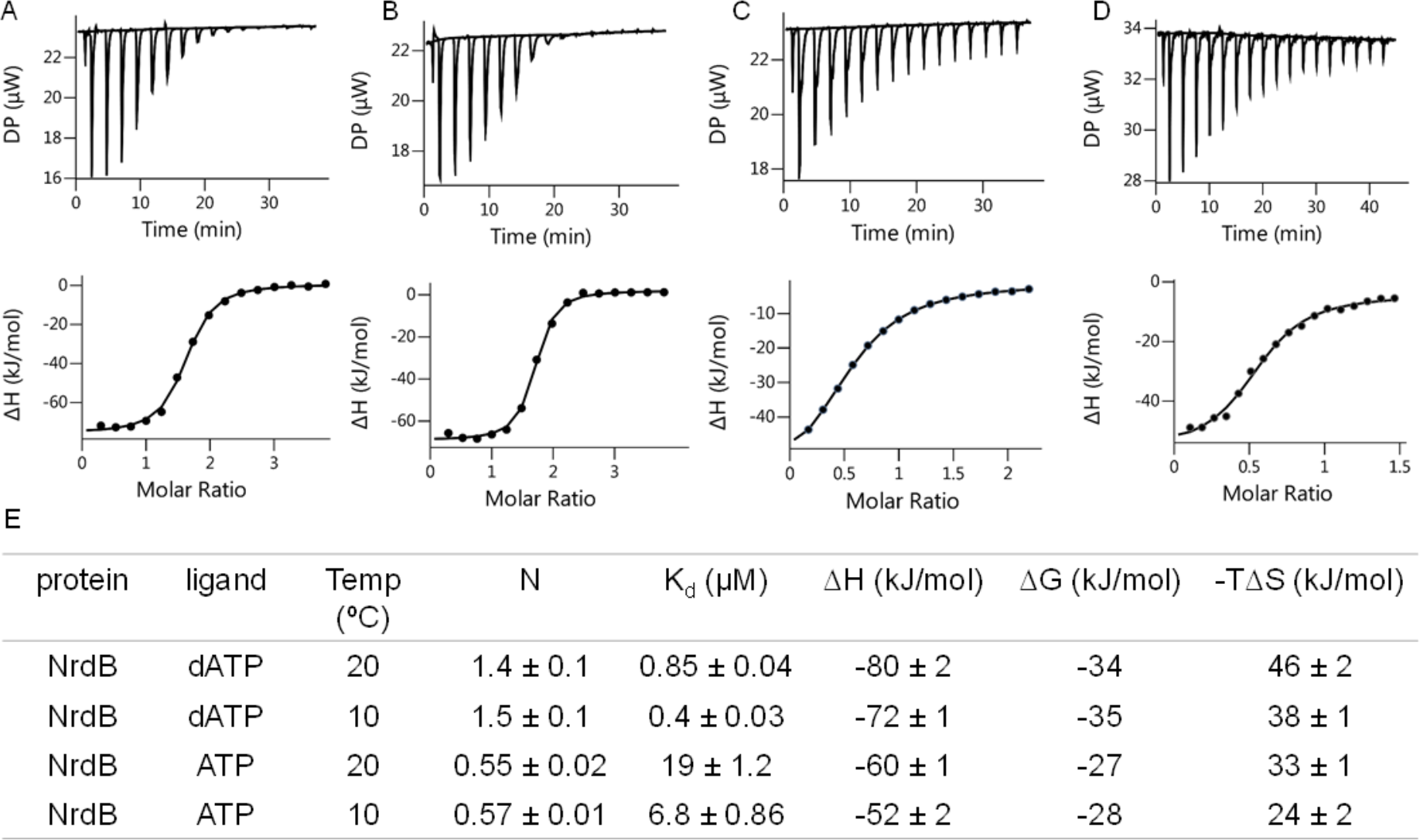
Representative ITC thermograms obtained by titration of ligands into NrdB. A) Titration of dATP to NrdB at 20°C, and B) titration of dATP to NrdB at 10°C. C) Titration of ATP to NrdB at 20°C, and D) titration of ATP to NrdB at 10°C. Isothermal calorimetric enthalpy changes are shown. E) Thermodynamic parameters of ligand binding to NrdB. Binding isotherms were fitted using a one-set-of sites binding model. All titrations were performed at 20°C and 10°C as described in Materials and methods.

We performed an additional set of ITC experiments at 10°C, which resulted in lower *K_d_* values (0.4 μM for dATP and 6.8 μM for ATP) but otherwise similar conclusions. Fitted stoichiometries were 1.5 ±0.1 for dATP and 0.57 ±0.01 for ATP in agreement with the 20°C results and underscoring our interpretation that the *F. ignava* NrdB protein binds two molecules of dATP and one molecule of ATP.

### Type of radical cofactor in the *F. ignava* NrdB protein

To elucidate the nature of the radical cofactor in the *F. ignava* NrdB protein we employed electron paramagnetic resonance (EPR) spectroscopy. X-band EPR spectra recorded on samples of NrdB∆169 expressed in the presence of excess Mn^2+^ and purified via affinity chromatography revealed an intense multiline signal with a signal width of 125 – 130 mT (Fig. 9). The signal varies in a uniform fashion in the interval 5 to 15 K and is thus attributed to a single paramagnetic species (Fig. 9, compare 5, 10 and 15 K spectra). Increasing the temperature further resulted in a complete disappearance of the signal at 30 K, with no new signal appearing. The shape, width and temperature dependence of the signal is in good agreement with an anti-ferromagnetically coupled Mn^III^/Mn^IV^ complex, where the complex line shape is a result of an S = ½ system where the unpaired electron is interacting with two I = 5/2 Mn centers. In a biological context, similar Mn^III^/Mn^IV^ species have been observed in the case of superoxidized Mn-catalase and as a short-lived intermediate during the assembly of the Mn^III^_2_-Y^•^ cofactor in NrdF (30,31). The presence of such an intense multiline signal in our purified samples suggests that this high-valent species is stable at least on an hours time-scale.

**Figure 9.**
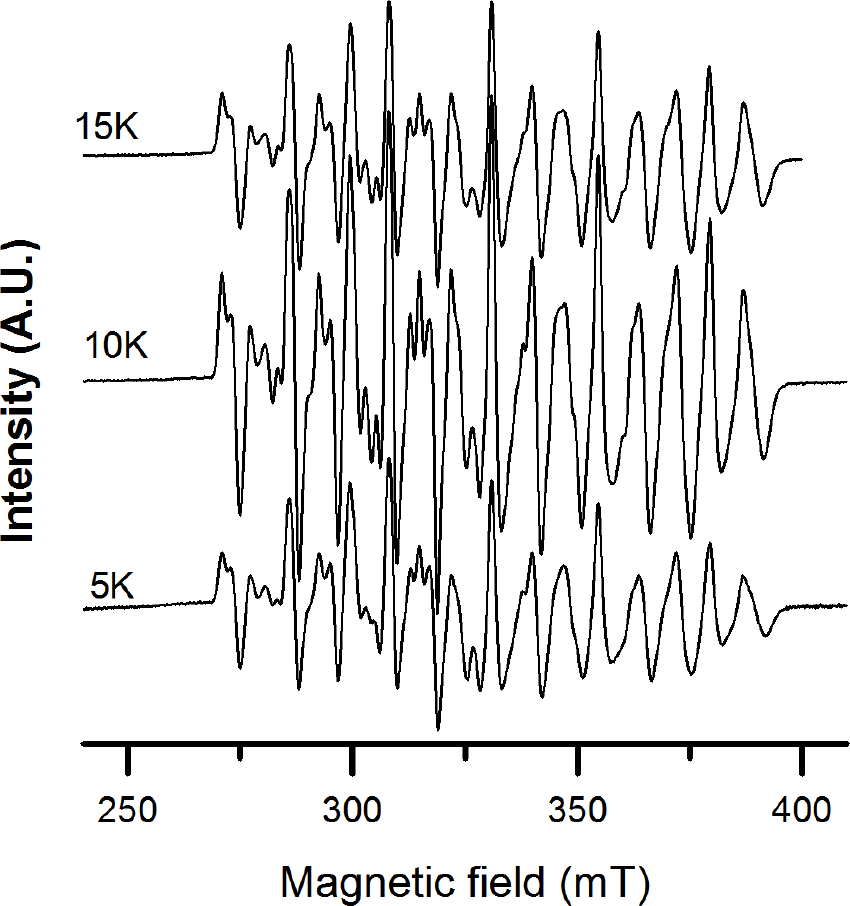
X-band EPR spectra of *F. ignava* NrdB∆169 recorded at 5 – 15 K. Samples recorded on 1.13 mM protein. Instrument settings: microwave frequency 9.28 GHz, modulation amplitude 10G, modulation frequency 100 kHz, microwave power 1 mW, temperature 5–15 K.

### RNR activity in mixtures of *F. ignava* and *L. blandensis* NrdA and NrdB proteins

The NrdB core of the firmicute *F. ignava* from residue 170 and onwards has extensive similarity (61% sequence identity, Fig. S5) to the core of the NrdB protein from the flavobacterium *L. blandensis*. They both harbor a mixed valent Mn_2_^III/IV^ center with capacity to initiate the radical-based enzyme reaction (this study and (8)). Both the *F. ignava* NrdB and the *L. blandensis* NrdB also harbor an ATP-cone domain that functions as an on/off switch for the activity of its RNR holoenzyme by forming tetrameric NrdB structures in presence of dATP to which the NrdA protein is prevented from binding in a productive fashion (above and (8)). However, the ATP-cones of *F. ignava* and *L. blandensis* NrdB proteins are more different (28% sequence identity, Fig. S5) and only aligns extensively over their C-terminal sequences, which in the *L. blandensis* structure has been shown to interact primarily with one of the two bound dATP molecules (8). The similarity between the two corresponding NrdA proteins is extensive (61% sequence identities, Fig. S6). Based on these similarities we designed a set of experiments to test whether RNR enzyme activity can be achieved in heterologous mixtures of *F. ignava* and *L. blandensis* NrdA and NrdB proteins and whether the unique Grx domain would disturb a heterologous interaction. Heterologous mixtures of class I RNR subunits have primarily been tested for distantly related enzymes, e.g. class I RNR subunits from *E. coli* and bacteriophage T4 with negative results (32). On the other hand, several thioredoxins are known to cross-react with heterologous RNRs, whereas Grxs usually do not (33).

Figure 10 shows that the heterologous *F. ignava* NrdA/*L. blandensis* NrdB holoenzyme is active and regulated by ATP and dATP via the ATP-cone linked to *L. blandensis* NrdB, whereas the heterologous *L. blandensis* NrdA/*F. ignava* NrdB holoenzyme is inactive. The same is true for heterologous mixtures with *F. ignava* NrdB∆Grx, as well as for *F. ignava* NrdB∆169 (Fig. 10A). *K_L_* for ATP is ≈300 μM and *K_i_* for dATP is ≈70 μM for the ATP-cone of *L. blandensis* NrdB in the heterologous mixture (Fig. 10B), i.e. more than 3-times higher than the *K_L(ATP)_* of 96 μM and the *K_i(dATP)_* of 20 μM for the *L. blandensis* holoenzyme (8). The *V_max_* obtained in the heterologous holoenzyme is 250 nmol/min•mg, corresponding to a *k_cat_* of approximately 0.2 s^−1^, approximately a fourth of the activity of the *L. blandensis* holoenzyme (Fig. 10A).

**Figure 10.**
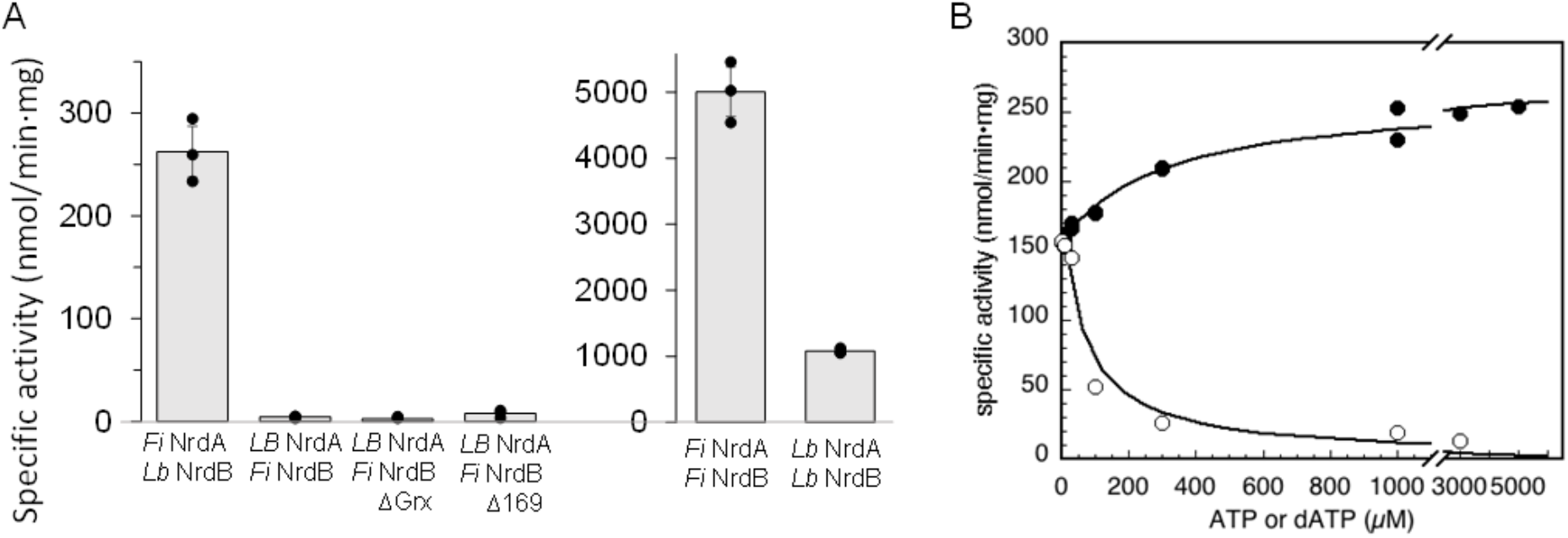
Enzyme activity in heterologous mixtures of NrdA and NrdB protein from *F. ignava* and *L. blandensis*. A) GDP reduction in presence of dTTP as effector. Assays were performed in triplicate with standard deviations shown. B) ATP (•) and dATP (○) titrations of *F. ignava* NrdA plus *L. blandensis* NrdB in presence of 2 mM dTTP and with GDP as substrate. ATP and dATP titrations in assays with CDP as substrate are shown in Fig. S7.

## Discussion

We have shown that the multidomain radical-generating component of the *F. ignava* class I RNR contains a gene fusion of an N-terminal Grx that is fully functional as a reductant of the RNR holoenzyme, and an ATP-cone that serves as a general on/off-switch of the enzyme. We also identified fusions of Grx-domains with NrdB proteins in *Francisella* spp., *Allofrancisella guangzhouensis* and several viruses (Fig. 1, Table S1), but none of the other cases were with the NrdBi subclass that the *F. ignava* protein belongs to. This strongly suggests that the *F. ignava grx::nrdBi* fusion was a separate evolutionary event, not related to the *grx::nrdB* fusions discovered in other organisms and viruses. On the contrary, the presence in *F. ignava* of a fusion between an ATP-cone domain and NrdBi appears to be the result of horizontal gene transfer since the majority of ATP-cones fused with *nrdBi* genes occur in flavobacteria (8). It thus appears most parsimonious to suggest that the ATP-cone::NrdBi fusion gene was first transferred to *F. ignava* and the gene was subsequently fused with the *grx* gene in the *F. ignava* genome.

Grx was first described as a physiological reductant for RNR in *Escherichia coli* (25), and has since also been observed to be involved in sulfate assimilation, detoxification, as well as development and proliferation, primarily in eukaryotic cells (26,34). Similarly to other redoxins, the active site of dithiol Grxs consists of a cysteine pair separated by two residues (predominantly -CPYC-) (26). The corresponding sequence in *F. ignava* Grx::NrdB is - CPWC- (Fig. S5). Grxs differ from other redoxins in that they form mixed disulfides with GSH and also promote glutathionylation/deglutathionylation reactions, which may lead to reduction of protein disulfides (34). *E. coli* Grx has been shown to use the dithiol mechanism in its reduction of *E. coli* RNR, whereas a human Grx was interpreted to reduce mammalian RNR via a glutathionylation mechanism (27). However, recent theoretical studies as well as thorough experimental studies on Grx-dependent reduction of protein disulfides with heterologous components from eukaryotic and bacterial origins show that the monothiol-dithiol mechanisms occur in parallel and that GSH concentration and dominance of specific steps in the mechanism determines the preferred path taken (35-37). In this study we show that the Grx::NrdB fusion protein of *F. ignava* can reduce its class I RNR holoenzyme via a dithiol mechanism, and that the C15S mutant in the Grx active site can reduce RNR less efficiently via a monothiol mechanism (Fig. S1). To our knowledge this is the first demonstration of parallel dithiol-monothiol reduction mechanisms in a native system between Grx and its oxidized substrate from the same species..

Over and above the fused Grx domain, the remarkable *F. ignava* NrdB protein exhibits two other unusual characters: a fused ATP-cone and a mixed valent Mn_2_^III/IV^ metal site. Both of these features have been recently observed in *L. blandensis* NrdB and in several NrdBi proteins in Flavobacteriales (8). The ATP-cone in *F. ignava* NrdB binds 2 dATP molecules like the cone in *L. blandensis* NrdB, but their amino acid sequences across the ATP-cones are surprisingly dissimilar in the N-terminal half (Fig. S5). This may relate to our finding that the cone in *F. ignava* binds only one ATP molecule, whereas the cone in *L. blandensis* binds 2 ATP-molecules. The ATP-loaded active *F. ignava* holoenzyme is α_2_β_2_, whereas the dATP-inhibited complexes are β_4_ for NrdB, and α_2_β_4_ plus α_4_β_4_ for the holoenzyme. All of these complexes were also observed in the *L. blandensis* RNR (8).

The mixed valent Mn_2_^III/IV^ metal site in *F. ignava* NrdB has a distinct EPR signal in the temperature range of 5-15K, with no other Mn-related EPR signals at 30K and no trace of a tyrosyl radical. A similar high valent Mn dimer was recently found to be present in NrdB from *L. blandensis* (8). Later, Boal and co-workers also reported a similar multiline signal in *Flavobacterium johnsoniae* class I RNR (9). However, in both of these latter cases the multiline feature represented only a fraction of the total metal content. Conversely, in our *F. ignava* NrdB samples presented here, the Mn_2_^III/IV^ signal is clearly the dominant metal species. These observations underscore the catalytic relevance of the Mn_2_^III/IV^ site, and support the notion that the NrdBi proteins represent a new subclass of class I RNRs, denoted subclass Id (6,8,9).

The similarities between *F. ignava* and *L. blandensis* NrdB proteins is further manifested by the enzyme activity observed in a heterologous mixture of *F. ignava* NrdA and *L. blandensis* NrdB, which is almost a third of that in the *L. blandensis* holoenzyme. Conversely, heterologous mixtures of *L. blandensis* NrdA and *F. ignava* NrdB lacks activity even in absence of the Grx domain or for the NrdB∆169 protein that lacks both the Grx domain and the ATP-cone. The *F. ignava* NrdB core may have undergone significant structural changes in order to accept the fusion of the Grx domain, which may also pertain to the divergent N-terminal of the ATP-cone sequence. Future studies will be directed to clarify this point.

All in all, we have shown that the unique NrdB protein in *F. ignava* carries an N-terminal Grx domain with capacity to act as a physiological reducant of its corresponding holoenzyme via a dithiol mechanism and less efficiently via a monothiol mechanism in the C15S mutant variant. The ATP-cone domain, which is fused between the Grx domain and the NrdB core functions as an allosteric on/off switch, promoting an enzymatically active α_2_β_2_ in presence of ATP and enzymatically inactive α_2_β_4_ and α_4_β_4_ complexes in presence of dATP. The radical cofactor in *F. ignava* NrdB is a mixed valent dinuclear Mn^III/IV^ site, which forms in the absence of an NrdI activase and lacks a tyrosyl radical. *F. ignava* NrdB is an enthralling illustration of how RNR subclasses continuously evolve via gain and loss of accessory domains and RNR-related proteins.

## Experimental procedures

### Bioinformatics

The RefSeq database (38) was downloaded 16 March 2018 and searched with the Pfam (39) profiles for Grx (PF00462) and the ATP-cone (PF03477) plus inhouse developed profiles for RNR proteins (http://rnrdb.pfitmap.org) using the HMMER software version 3.1b2 (40). For RNR proteins, only hits covering at least 90% of the length of the profile were kept. For the Grx and ATP-cone profiles, only hits with a higher bitscore than the Pfam specified GA scores (21.50 in both cases) were kept.

### Cloning

DNA fragments encoding NrdAi (WP_006702002) and NrdBi (EKB53615/WP_006702003) were amplified by PCR from Facklamia ignava CCUG 37419 genomic DNA, obtained from the Culture Collection, University of Gothenburg using specific primers: NrdA: FiR1_For 5’-tctcCATATGACCGCACAATTAAAGAATC-3’ and FiR1_Rev 5’- cagaGGATCCTTAAGCTTCACAAGCTAAGC -3’. NrdB: FiR2_For 5’- tctaCATATGACTCAAGTACAAGTTTATAG-3’, FiR2_REV 5’- cagaGGATCCTTAGAATAGGTCGTCGGC-3’, The PCR products were purified, cleaved with NdeI and BamHI restriction enzymes and inserted into a pET-28a(+) expression vector (Novagen, Madison, Wisconsin, USA). The obtained constructs pET-nrdA and pET-nrdB contained an N-terminal hexahistidine (His) tag and thrombin cleavage site. To construct the truncated NrdB mutant, lacking the Grx domain (residues 1-78), new forward primer FiR2∆Grx_For 5’- tctaCATATGAGCAAAATCCCGCAACAC -3’ was used with FiR2_REV to yield a pET-nrdB∆Grx. To construct the truncated NrdB mutant, lacking both the Grx and the entire ATP-cone domains (residues 1-169), new forward primer FiR2∆169_For 5’- tctaCATATGGCGCGTCAACGTGATATA -3’ was used with FiR2_REV to yield a pET- nrdB∆169. The cloning process and the resulting constructs pET-nrdB∆Grx and pET- nrdB∆169 were similar to that of the wild type pET-nrdB, except that they lacked sequences coding for the N-terminal 78 and 169 amino acids respectively. To obtain NrdB bearing point mutations of individual cysteine residues to serines at the Grx active site, pET-nrdB_C12S, pET-nrdB_C15S or the double mutant, were both cysteines were mutated to serines pET- nrdB_C12SC15S, constructs containing nucleotide mismatches T34A, G44C and T34AG44C, were ordered from GenScript.

### Protein expression

Overnight cultures of *E. coli* BL21(DE3)/pET28a(+) bearing pET-nrdA, pET-nrdB, pET- nrdB∆Grx, pET-nrdB∆169, pET-nrdB_C12S, pET-nrdB_C15S, pET-nrdB_C12SC15S were diluted to an absorbance at 600 nm of 0.1 in LB (Luria-Bertani) liquid medium, containing kanamycin (50 μg/ml) and shaken vigorously at 37°C. At an absorbance of 0.8 A_600_ isopropyl-β-D-thiogalactopyranoside (Sigma) was added to a final concentration of 0.5 mM; the cultures expressing NrdB were further supplementented with MnSO_4_ (final concentration 0.5 mM) during the induction. The cells were grown overnight at 30°C and harvested by centrifugation.

### Protein purification

The cell pellet was resuspended in lysis buffer: 50 mM Tris-HCl pH 7.6 containing 300 mM NaCl, 10% glycerol, 2 mM DTT, 10 mM imidazole, 1 mM PMSF. Cells were disrupted by high pressure homogenization and the lysate was centrifuged at 18,000 × g for 45 min at 4°C.

The recombinant His-tagged protein was first isolated by metal-chelate affinity chromatography using ÄKTA prime system (GE Healthcare): the supernatant was loaded on a HisTrap FF Ni Sepharose column (GE Healthcare), equilibrated with lysis buffer (without PMSF), washed thoroughly with buffer and eluted with buffer containing 500 mM imidazole. NrdB_C12S, NrdB_C15S, NrdB_C12SC15S and the wild type NrdB used for measuring the redox activity of the NrdB fused Grx were then applied to the Sephadex G-25 PD10 desalting column equilibrated with buffer containing 50 mM Tris-HCl pH 7.6, 300 mM NaCl, 10% glycerol and 1 mM DTT, frozen in liquid nitrogen and stored at −80°C until used.

For NrdA, NrdB, NrdB∆Grx and NrdB∆169 further purification was accomplished by fast protein liquid chromatography (FPLC) on a 125 ml column packed with HiLoad 16/600 Superdex 200 pg column (GE Healthcare) using ÄKTA prime system, equilibrated with buffer containing 50 mM Tris-HCl pH 7.6, 300 mM NaCl, 10% glycerol and 2 mM DTT. Eluted protein was collected.

NrdA was further applied to HIC chromatography using the HiLoad 16/60 phenyl sepharose column (GE Healthcare) in 50 mM Tris-HCl pH 7.6, 2 mM DTT, 0.75 M (NH_4_)_2_SO_4_, washed extensively (15 column volumes) with the same buffer and eluted with buffer without ammonium sulphate. The protein was resuspended in excess of buffer containing 50 mM Tris-HCl pH 7.6, 300 mM NaCl, 10% glycerol, 2 mM DTT, concentrated and frozen until used. The HIC chromatography removed residual nucleotide contamination from NrdA.

*L. blandensis* NrdA and NrdB were expressed and purified as previously described (8)

Protein concentrations were determined by measuring the UV absorbance at 280 nm based on protein theoretical extinction coefficients 99 700 M^−1^ cm^−1^ for NrdA, 72 770 M^−1^ cm^−1^ for NrdB (and cysteine to serine mutants), 54 320 M^−1^ cm^−1^ for NrdB∆Grx and 51 340 M^−1^ cm^−1^ for NrdB∆169. Protein purity was evaluated by SDS–PAGE (12%) stained with Coomassie Brilliant Blue. Proteins were concentrated using Amicon Ultra-15 centrifugal filter units (Millipore), frozen in liquid nitrogen and stored at −80°C until used.

For EPR measurements, NrdB∆169, was purified using affinity chromatography as described above but transferred to EPR tubes and flash frozen in liquid nitrogen in EPR tubes immediately upon elution.

### Enzyme activity measurements

Enzyme assays were performed at room temperature in 50 mM Tris-HCl, pH 8 in volumes of 50 μl. Reaction conditions, giving maximal activity were determined experimentally. In a standard reaction the constituents were; 10 mM DTT, 40 mM or 20 mM Mg(CH_3_CO_2_)_2_ (when NrdA of *F. ignava* or *L.blandensis* was used respectively), 10 mM KCl, 0.8 mM CDP, and various concentrations of allosteric effectors ATP or dATP. Mixtures of 0.1 μM to 1 μM of NrdB, 0.07 μM NrdB∆169, 0.5 μM NrdB_C12S, NrdB_C15S, or NrdB_C12SC15S and a 10- fold excess of NrdA were used. Some components were explicitly varied as indicated in specific experiments.

In experiments aimed to determine the redoxin activity of the NrdB fused Grx, DTT was omitted. Instead, 4 mM or 10 mM reduced GSH, 11 μg ml^−1^ glutathione reductase (from yeast, Sigma) and 1 mM NADPH were added to the reaction mixtures. CDP (0.8 mM) was used as substrate and ATP (3 mM) as effector. Protein concentration of 0.5 μM for wild type NrdB, NrdB_C12S, NrdB_C15S, or NrdB_C12SC15S were used in combination with 5 μM NrdA.

When dTTP (2 mM) was used as an s-site effector, 0.8 mM GDP was used as substrate. In the four substrate assays, the substrates CDP, ADP, GDP and UDP were simultaneously present in the mixture at concentrations of 0.5 mM each with 2 mM of one of the effectors (ATP, dTTP or dGTP). The substrate mixture was added last to start the reactions.

Enzyme reactions were incubated for 2 to 30 minutes at room temperature and then stopped by the addition of methanol. Substrate conversion was analyzed by HPLC using a Waters Symmetry C18 column (150 × 4.6 mm, 3.5 μm pore size) equilibrated with buffer A. Samples of 25-100 μl were injected and eluted at 0.4 ml/min at 10°C with a linear gradient of 0–30% buffer B over 40 min for the separation of CDP and dCDP or 0-100% buffer B over 45 min for the separation of GDP and dGDP (buffer A: 10% methanol in 50 mM potassium phosphate buffer, pH 7.0, supplemented with 10 mM tributylammonium hydroxide; buffer B: 30% methanol in 50 mM potassium phosphate buffer, pH 7.0, supplemented with 10 mM tributylammonium hydroxide). Compound identification was achieved by comparison with injected standards. Relative quantification was obtained by peak height measurements in the chromatogram (UV absorbance at 271 nm or 254 nm) in relation to standards. Specific activities are given as nmol product formed per min and mg protein.

From a direct plot of activity versus concentration of effector, the K_L_ values for binding of effectors to the s-site and the a-site, were calculated in SigmaPlot using the equation:

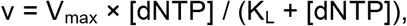

and K_i_ for non-competitive dATP inhibition at NrdB was calculated in Sigmaplot using the equation:

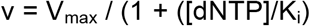

### Photometric activity assays

Photometric assays for NrdB-fused Grx based on the artificial electron acceptor HED were performed as described in earlier studies (41,42). The standard Grx assay contains 50 mM Tris pH 8.0, 0.1 mg/ml BSA, 11 μg ml^−1^ glutathione reductase (from *S. cerevisiae*), 4 mM glutathione, 0.75 mM HED, and 0.4 mM NADPH. The above ingredients were mixed and incubated for 3 minutes, after which the reaction was started by the addition of 0.1 μM wild type or mutant Grx (NrdB fused). Reference cuvette contained all ingredients, except Grx. A_340_ was recorded for 3 min at room temperature using Lambda 35 UV/VIS spectrophotometer (Perkin Elmers). Linear decrease in A_340_ was used to calculate moles of NADPH consumed using its extinction coefficient of 6220 M^−1^ cm^−1^.

The combined redoxin/RNR assays contained 0.5 μM NrdB, 5 μM NrdA, indicated amount of glutathione, 11 μg ml^−1^ glutathione reductase, 0.25 mM NADPH, 10 mM Mg(CH_3_CO_2_)_2_, 3 mM ATP. Reaction was started by the addition of 0.8 mM CDP. The reaction was monitored by the change of A_340_ using Cary 60 UV-VIS spectrophotometer (Agilent technologies) Specific activity was calculated using 1 mol of NADPH equals 1 mol of formed dCDP.

### GEMMA analysis

In GEMMA, biomolecules are electrosprayed into gas phase, neutralized to singly charged particles, and the gas phase electrophoretic mobility is measured with a differential mobility analyzer. The mobility of an analyzed particle is proportional to its diameter, which therefore allows for quantitative analysis of the different particle sizes contained in a sample (43). The GEMMA instrumental setup and general procedures were as described previously (44). NrdA, NrdB, NrdB∆Grx and NrdB∆169 proteins were equilibrated by Sephadex G-25 chromatography into a buffer containing 100 mM NH_4_CH_3_CO_2_, pH 7.8 and 2 mM DTT. Prior to GEMMA analysis, the protein samples were diluted to a concentration of 0.05 mg/ml in a buffer containing 20 mM NH_4_CH_3_CO_2_, pH 7.8, 1 mM DTT, 0.005% (v/v) Tween 20, nucleotides (when indicated), and Mg(CH_3_CO_2_)_2_ (equimolar to the total nucleotide concentration), incubated for 5 min at room temperature, centrifuged and applied to the GEMMA instrument. The runs were conducted at low flow rate, resulting in 1.4 – 2 Psi pressure. The GEMMA system contained the following components: 3480 electrospray aerosol generator, 3080 electrostatic classifier, 3085 differential mobility analyzer, and 3025A ultrafine condensation particle counter (TSI Corp., Shoreview, MN).

### Analytical SEC

The SEC experiments were performed at ambient temperature with a Superdex^TM^ 200 10/300 column (GE Healthcare) equilibrated with a mobile phase containing 50 mM KCl, 10 mM MgCl_2_ 0.1 mM dithiothreitol, and 50 mM Tris-HCl pH 7.6. 3 mM ATP or 0.1 mM dATP was also included in the mobile phase when nucleotide-dependent protein oligomerization was studied. The injection loop volume was 100 μl and the flow rate was 0.5 ml/min. The UV trace was recorded with a Jasco UV-2075 Plus detector (Jasco Inc., Easton, MD) at 290 nm to limit the absorbance from the nucleotides, The proteins were incubated in mobile phase for 5 min prior to injection onto the column.

### Isothermal titration calorimetry measurements

Isothermal titration calorimetry (ITC) experiments were carried out on a MicroCal ITC 200 system (Malvern Instruments Ltd) in a buffer containing 50 mM Tris pH 7.65, 300 mM NaCl, 10% glycerol, 2 mM tris(2-carboxyethyl)phosphine, and 10 mM MgCl_2_. Measurements were done at 20°C and 10°C. The initial injection volume was 0.5 μl over a duration of 1 s. All subsequent injection volumes were 2-2.5 μl over 4-5 s with a spacing of 150 −180 s between the injections. Data for the initial injection were not considered. For dATP binding analysis, the concentration of NrdB in the cell was 40 μM and dATP in syringe 600 μM. For titration of ATP into NrdB, cell and syringe concentrations were 103 μM NrdB and 1.2 mM ATP. The data were analyzed using the built-in one set of sites model of the MicroCal PEAQ-ITC Analysis Software (Malvern Panalytical). Standard deviations in thermodynamic parameters, N and *K_d_* were estimated from the fits of three different titrations.

### Electron paramagnetic resonance spectroscopy

Measurements were performed on a Bruker ELEXYS E500 spectrometer using an ER049X SuperX microwave bridge, in a Bruker SHQ0601 cavity equipped with an Oxford Instruments continuous flow cryostat, and using an ITC 503 temperature controller (Oxford Instruments). The Xepr software package (Bruker) was used for data acquisition and processing of spectra.

## Acknowledgments

We thank Al Claiborne, Wake Forest University, NC, USA, for useful advice on bacillithiol and glutathione biosynthesis, Ann Magnusson and Sigrid Berglund, Uppsala University, Sweden, for valuable discussions on dinuclear manganese centers, Ilya Borovok, Tel-Aviv University, Israel, for discussions on ATP-cones, and Malvern Panalytical for kindly sharing the MicroCal PEAQ-ITC Analysis Software for the analysis of ITC data. This study was supported by grants from the Swedish Cancer Society (CAN 2016/670 to BMS), the Swedish Research Council (2016–01920 to BMS), the Wenner-Gren Foundations (to BMS) and the Carl Trygger Foundation (to AH). Work in the laboratory of GB is supported by the Swedish Research Council (621–2014-5670), the Swedish Research Council for Environment, Agricultural Sciences and Spatial Planning (213-2014-880) and the European Research Council (714102).

## Abbreviations and nomenclature

a-site: allosteric overall activity site in the ATP-cone
BSA: bovine serum albumin
dNTPs: deoxyribonucleotides
DTT: dithiothreitol
EPR: electron paramagnetic resonance
FPLC: fast protein liquid chromatography
GEMMA: gas-phase electrophoretic macromolecule analysis
Grx: glutaredoxin
GSH: glutathione
HED: 2-hydroxyethyl disulfide
HIC: hydrophobic interaction chromatography
HPLC: high performance liquid chromatography
ITC: isothermal titration calorimetry
NrdB∆Grx: NrdB protein lacking 69 N-terminal residues corresponding to the glutaredoxin domain
NrdB∆169: NrdB protein lacking the glutaredoxin domain and the ATP-cone domain
PCR: polymerase chain reaction
PMSF: phenylmethylsulfonyl fluoride
RNR: ribonucleotide reductase
SDS-PAGE: sodium dodecyl sulphate polyacrylamide gel electrophoresis
SEC: size exclusion chromatography
s-site: allosteric specificity site in NrdA

